# Interactions of surfactants with the bacterial cell wall and inner membrane: Revealing the link between aggregation and antimicrobial activity

**DOI:** 10.1101/2021.05.26.445833

**Authors:** Pradyumn Sharma, Rakesh K. Vaiwala, Srividhya Parthasarathi, Nivedita Patil, Morris Waskar, Janhavi S. Raut, Jaydeep K. Basu, K. Ganapathy Ayappa

## Abstract

Surfactants with their intrinsic ability to solubilize lipids are widely used as antibacterial agents. Interaction of surfactants with the bacterial cell envelope is complicated due to their propensity to aggregate. It is important to discern the interactions of micellar aggregates and single surfactants on the various components of the cell envelope to improve selectivity and augment the efficacy of surfactant-based products. In this study, we present a combined experimental and molecular dynamics investigation to unravel the molecular basis for the superior kill efficacy of laurate over oleate observed in contact time assays with live *E. coli*. To gain a molecular understanding of these differences, we performed all-atom molecular dynamics simulations to observe the interactions of surfactants with the periplasmic peptidoglycan layer and the inner membrane of Gram-negative bacteria. The peptidoglycan layer allows a greater number of translocation events for laurate when compared with oleate molecules. More interestingly, aggregates did not translocate the peptidoglycan layer, thereby revealing an intrinsic sieving property of the bacterial cell wall to effectively modulate the surfactant concentration at the inner membrane. The molecular dynamics simulations exhibit greater thinning of the inner membrane in the presence of laurate when compared with oleate, and laurate induced greater disorder and decreased the bending modulus of the inner membrane to a greater extent. The enhanced antimicrobial efficacy of laurate over oleate was further verified by experiments with giant unilamellar vesicles, which revealed that laurate induced vesicle rupture at lower concentrations in contrast to oleate. The novel molecular insights gained from our study uncovers hitherto unexplored pathways to rationalize the development of antimicrobial formulations and therapeutics.

## Introduction

Surfactants and fatty acids with their ability to solubilize lipid membranes are one of the earliest known antimicrobials used widely due to their broad spectrum activity against bacteria, viruses, and fungi ^1,2^. Bio-surfactants are also emerging as alternatives to synthetic surfactants due to their low toxicity and biodegradability ^3^. Since the common building blocks of both microbial and mammalian cell membranes ^4^ are phospholipids, surfactants can lyse a wide class of cellular systems. Given the complexity of the bacterial cell envelope and the differences between Gram-negative and Gram-positive cell architectures, a molecular understanding of surfactant interactions with the bacterial cell envelope is needed to improve selectivity and augment the efficacy of antibacterial action. Additionally, understanding the inhibitory mechanisms of surfactants at molecular scales is vital for adequately assessing the scope and extent of various formulations used as disinfectants and delivering maximum hygiene benefits.

The cell envelope of Gram-negative bacteria has an outer membrane (OM) made up of lipopolysaccharides and lipids, an intervening periplasmic peptidoglycan (PGN) layer, and a phospholipid inner membrane (IM)^5^, while Gram-positive bacteria are characterized by a thick peptidoglycan layer and an inner membrane. Surfactants and other antimicrobials first bind to the OM of the bacterial cell envelope and penetrate the bacterial cell wall prior to interacting and solubilizing the phospholipid inner membrane. Due to the negatively charged cell surfaces of both Gramnegative and Gram-positive bacteria, cationic surfactants have been widely used as antibacterial cleansing agents. In particular, quaternary ammonium surfactants ^1,6^ and fatty acids ^7–9^ have been extensively investigated. The binding efficacy of the surfactant is a function of several factors which include size, charge, molecular architecture, and collective properties such as the critical micellar concentration and aggregation numbers^10^ which differ based on their protonation states ^11,12^, chain length, and extent of saturation. For non-ionic surfactants like disaccharide monoesters, the carbon chain length has been perceived to be the most crucial factor influencing their antimicrobial activity ^13^. N-acyl surfactants show variation in antibacterial properties based on chain length and degree of unsaturation ^14^ with the presence of double bonds intensifying the antibacterial activity. Hence a surfactant specific mechanism is anticipated for these molecules based on chain length, the extent of saturation, charge, and concentration.

Despite the wide use of surfactants and fatty acids as antimicrobial agents, the molecular interactions of these molecules with various components of the bacterial cell envelope are incompletely understood largely due to the inherent complexity of the cell envelope. Whether surfactants solubilize the OM or penetrate the OM through channels to access the IM are open questions. Recent in vitro studies on model bacterial membrane platforms, coupled with super resolution microscopy methods indicate that model membrane constructs can potentially be used to assess the barrier characteristics of bacterial membranes^15–17^. However, these experiments are challenging both from the point of constructing the reliable bacterial cell wall mimics and using appropriate microscopic tools to interrogate the membrane in the presence of external agents^15,18^.

In recent years, molecular dynamics (MD) simulations have evolved as a powerful tool to study bacterial membranes and assess their interactions and free energy barriers with small molecules and antibiotics. MD simulations have provided a molecular understanding of the barriers offered by different regions of the complex OM^15,19^ highlighting the asymmetric free energy landscape for molecule translocation^19^ which is quite distinct from the IM^15^. The barrier properties of the PGN layer have only recently been investigated in our laboratory^20^. Molecular dynamics simulations of surfactants and fatty acids have been widely used to capture properties like self-assembly and partitioning of surfactants^21,22^ and also used for investigating the interactions of surfactants with mammalian membrane models^23,24^. However, owing to the complex architecture of the bacterial cell envelope, MD simulations of interactions with surfactants are yet to be reported. Antibacterial properties of surfactants, the focus of this manuscript, which are salts of the fatty acids, have been studied to a lesser extent, and although their antimicrobial properties are known, a molecular view of their interactions with the cell envelope and subsequent action is only partially understood.

In this study, we investigate the antimicrobial properties of sodium salts of laurate and oleate and perform atomistic MD simulations to provide a molecular explanation for the increased bacterial kill efficacy of laurate over oleate as observed in contact time kill experiments carried out with *E. coli*. The novelty of our analysis lies in studying the interaction of surfactants with the PGN layer in addition to the phospholipid IM. Simulations with different surfactant concentrations allow us to study the influence of aggregation behaviour on the passage of surfactant molecules through PGN and also assess the interactions with peptide and glycan moieties with PGN. A detailed analysis to study the influence of surfactants on the IM properties such as membrane thickness, in-plane lipid order, and bending modulus in the presence of surfactants is carried out. Combined with rupture data from giant unilamellar vesicles, we attribute the contrasting efficacies of laurate and oleate to the differences in chain length dependent aggregation behavior of these molecules, which plays a pivotal role in the barrier offered to them by PGN. Our study also shows that once surfactants partition into the IM, the extent to which they perturb the membrane is a function of the surfactant chain length and concentration.

## Materials and Methods

### Bacteria kill assays

Surfactants used in these studies (sodium oleate & sodium laurate) were procured from Sigma Aldrich. The test bacteria *E. coli* procured from American Type Culture Collection (ATCC 10536) were grown overnight on Tryptic Soy agar (TSA) plate (procured from Difco ™) at 37°C and incubated for 16 hours. The cell density was adjusted at optical density 620 (OD_620_) using a spectrophotometer to get the final count of 10^8^ cfu/mL for *E. coli*. 1 mL of the bacterial suspension was added to 9 mL sterile distilled water containing the test material. After a contact time of 5 min with respective concentrations of sodium oleate and sodium laurate, 1 mL of the sample was withdrawn and added to 9 mL of D/E neutralizing broth purchased from Difco ™. The residual bacteria were enumerated by serial dilution of the sample and plating it using TSA. After solidification, these plates were incubated at 37°C for 24 hrs. The colonies on the plates were counted after 24 hrs, and log reduction is calculated by comparing with the culture control. For the kill kinetics study, the contact time of bacteria with 40 mM sodium oleate and 40 mM sodium laurate was varied from 1 min, 2 mins, 5 mins, and 10 mins, followed by neutralization, dilutions, and plating.

### Molecular Dynamics Simulations

All-atom simulations were performed using GROMACS version 5.1.4^25^. The PGN model was taken from our previous study^20^ where a CHARMM36 compatible forcefield is used^26^. Surfactant-incorporated IM models were obtained from CHARMM-GUI web server^27^. Inner membrane model was procured from our previous work^15^. The membranes containing laurate and oleate have been studied at surfactant molar concentrations 20% and 40%, as summarized in Table S2. In all the membranes studied, the lipid compositions for DOPE:DOPG:TOCL (TOCL1) correspond to the inner membrane of *E. coli*, viz. ∼75:20:5. CHARMM36 force-field^28^ was used for surfactants, lipids, and ions, while modified TIP3P^29^ water model was employed to model aqueous solvent. The potassium ions (K^+^) were added to maintain electroneutrality. Simulation details for all the systems examined are given in the SI.

### GUV preparation and fixing

Giant unilamellar vesicles (GUVs) used in the experiments were prepared by the electroformation method. The GUVs were attached to the DPPC bilayer through biotin-streptavidin bonds using a similar protocol to our earlier studies^30^. GUVs fixed to the glass substrate were imaged using a Leica TCS SP5 microscope. The detailed protocol is described in the SI.

## Results

### Efficacy of shorter surfactants

We have studied the dose-dependent effect of sodium laurate and sodium oleate on *E. coli* viability using contact time kill (CTK) assays (Fig. 1A) at room temperature (25°C) and a pH value of 8-8.5. A small decrease in the viable population of bacteria for both surfactants at lower concentrations (< 10 mM) is observed. At 20 mM, about a 2 log order decrease is observed for laurate, and above 20 mM complete kill of bacteria was observed only in the case of laurate with oleate showing minimal antibacterial activity. Fig. 1B illustrates the temporal evolution of the population for a 40 mM surfactant concentration, indicating a complete kill with laurate at 5 min with about 1 log order reduction in the case of oleate. In order to determine the origins of these differences in activity, we study the interaction of laurate and oleate with PGN and IM using MD simulations and GUV experiments.

**Fig. 1.**
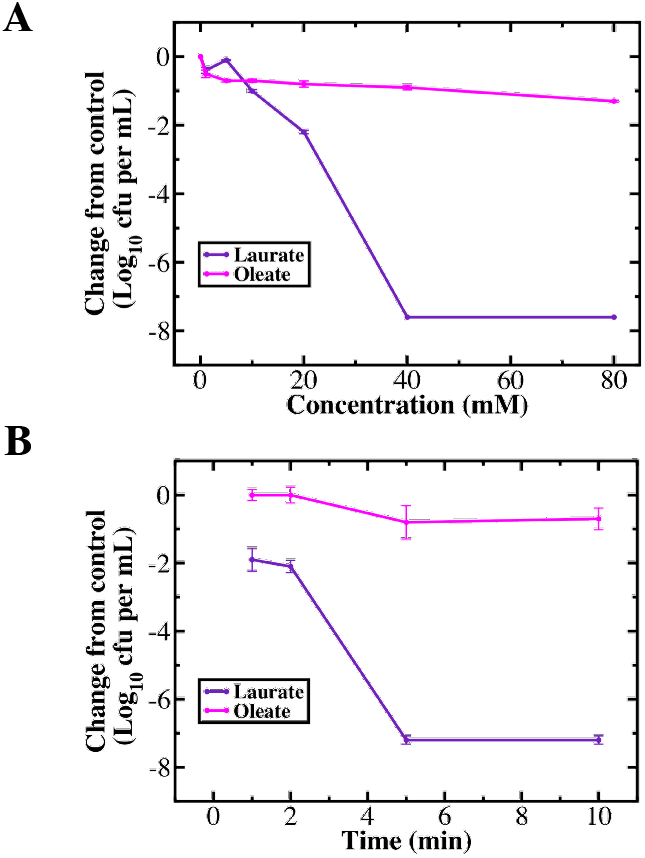
(A) Dose-dependent effects of sodium laurate and sodium oleate on *E. coli* viability. (B) *E. coli* kill kinetics with sodium laurate and sodium oleate. Change from control is defined as log_10_ (Test) - log_10_ (Control). A distinct increase in kill efficacy is observed for sodium laurate when compared with sodium oleate.

### Peptidoglycan - The unexplored barrier

We first examined the barrier properties of the PGN layer to surfactant molecules. Initially, we studied the interactions of these surfactants with PGN utilizing all-atom molecular dynamics simulations of a single surfactant molecule with the PGN layer of Gram-negative bacteria (Table S1). Translocation events in all systems were based on the center of mass trajectories of the PGN layer and the surfactant molecules as described in the SI (Fig. S1). Each of the systems with a single laurate (L1) and oleate (O1) was simulated for 500 ns, and multiple translocation events occurred for both L1 and O1 (Figs. 2A, B, and C), providing explicit confirmation for the lack of any significant barrier in the PGN layer for a single surfactant molecule. Interestingly, we have also observed differences in the number of translocation events for a single molecule of laurate and oleate during the 500 ns duration (Fig. 2C). For laurate, 13 translocation events were observed, while for oleate, only five such events occurred, and representative trajectories illustrate that laurate rapidly crosses the PGN layer (Fig. 2A), while oleate resides for about 8 ns (Fig. 2B) in the vicinity of the PGN layer. Similar trends were observed for several such events, affirming that PGN has a stronger interaction with oleate when compared with laurate, resulting in higher translocation events for the latter (Fig. 2C).

**Fig. 2.**
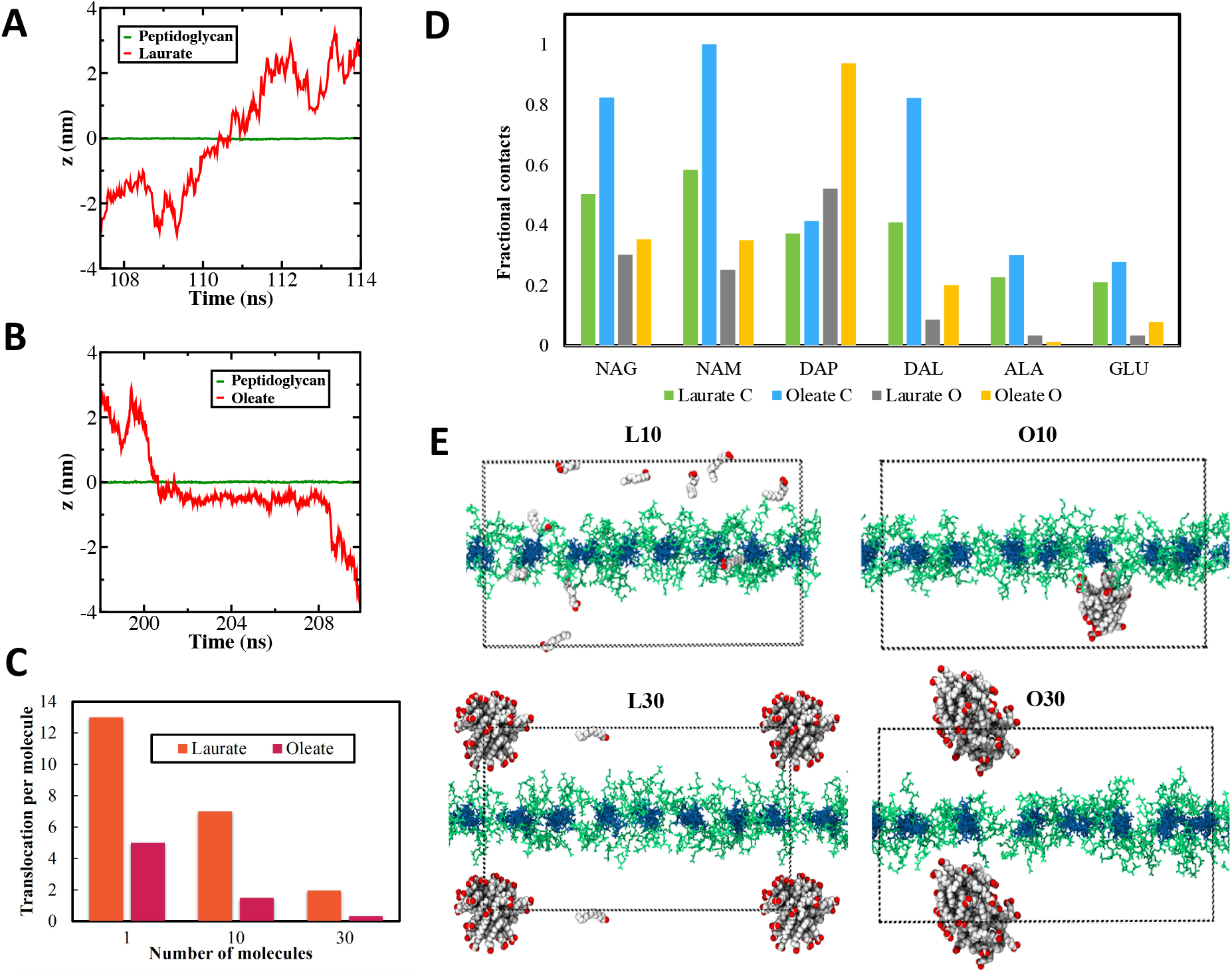
Time evolution of the center of mass of z-coordinates of the (A) laurate (red), (B) oleate (red), and the PGN (green) layer. (C) The number of translocation events for surfactant molecules during 500 ns simulation in all the PGN systems. (D) The fractional contacts made by carbon and oxygen atoms of surfactants with the peptidoglycan subunits during the course of a single surfactant simulations (L1 and O1). We have used a 0.4 nm cut-off to define contact and calculated these numbers for 500 ns simulations. The numbers of contacts for carbon atoms are normalized with respect to the number of carbon atoms in each surfactant molecule. Reported numbers are further scaled by the highest contacts (837) obtained in the case of oleate carbon atoms and NAM. (E) Snapshots for systems containing 10 and 30 molecules of laurate (L10 and L30) and oleate (O10 and O30) at the end of 500 ns simulations.

To understand the molecular details of these trends, we have calculated the fractional contacts Fig. 2D) between carbon (C) and oxygen (O) atoms of surfactant molecules with the peptidoglycan subunits, namely N-acetylglucosamine (NAG), N-acetylmuramic acid (NAM), L-alanine (ALA), D-iso-glutamate (GLU), meso-diamino pimelic acid (DAP), and D-alanine (DAL). We have employed a cutoff of 0.4 nm to define these contacts based on the center-of-mass coordinates. Surfactant carbon atom contacts are greater when compared with the oxygen head-groups for all peptidoglycan subunits except DAP. The cationic site present in DAP causes a preferential electrostatic attraction for the anionic headgroup of the surfactants. With the exception of ALA, oleate headgroups have a greater number of contacts when compared with the laurate headgroups. In general, the increased number of contacts for oleate result in greater residence times for oleate in the vicinity of the peptidoglycan layer (Fig. 2B). It can be summarized that the differences in the number of translocation events for the two surfactants are determined by the interplay between the interactions of surfactant molecules with sugars and amino acids present in the PGN layer.

To observe interactions at higher concentrations, 500 ns simulations with 10 (L10 and O10) and 30 (L30 and O30) surfactant molecules (Table S1) were performed by initially placing surfactants randomly on either side of the PGN layer. We observed surfactant aggregation for O10, O30, and L30 systems, although no aggregate was formed for L10 (Fig. 2E) over the course of the simulation. The differences in the tendency for aggregate formation can be attributed to the different critical micellar concentrations (CMC) for these surfactants, as observed in previous molecular dynamics studies^22^. The systems having 10 and 30 molecules of surfactants correspond to 11 mM and 37 mM surfactant concentrations, respectively. Potassium oleate having a lower CMC value of 1 mM, formed a micellar aggregate in the O10 system (Fig. 2E), while potassium laurate with a higher CMC value of 25 mM did not form the aggregate in L10 (Fig. 2E). The aggregate formation was however observed for both L30 and O30 systems. Interestingly these aggregates were unable to translocate through the PGN layer, resulting in lower translocation events at higher concentrations (Fig. 2C). Oleate has shown a lower number of events (Fig. 2C), similar to the single-molecule simulations.

Furthermore, to understand the difference in aggregation time scales, we have performed a cluster analysis (Fig. S2). In this analysis, one surfactant molecule is considered as an individual cluster in itself, and once a molecule comes into the vicinity (0.35 nm) of another molecule or aggregate, it becomes part of the corresponding cluster or aggregate. Hence a decreasing trend in the number of clusters denotes the formation of larger aggregates. Oleate systems O10 and O30 formed large aggregates within 200 ns due to lower CMC value. Laurate system L30 has shown contrasting results, with a single molecule remaining isolated even after 500 ns (Fig. 2E). The fast aggregation kinetics of oleate compared to laurate, coupled with the inability of aggregates to cross the PGN layer, results in a further decline in the translocation events for oleate (Fig. 2C).

### Surfactant induces thinning in bacterial inner membranes

We further studied the structural and mechanical properties of the surfactant incorporated Gram-negative IM having 20% and 40% surfactants using 1 *µ*s atomistic simulations (Fig. S3 and Table S2). The density of lipid molecules was computed along the z-direction, normal to the membrane plane, and we observe an overall decrease in the DOPE density upon addition of surfactant, giving rise to a more uniform density variation within the membrane (Fig. 3A). The intensity of the well defined peaks in the vicinity of the headgroups observed for IM decrease with increasing surfactant concentration. Similar trends were observed for the DOPG and TOCL density distributions illustrated in Fig. S4. This provides the first signatures of surfactant induced disruption of lipid packing within the bilayer. Although a decrease in the overall bilayer density is observed, the density variation in the bilayer mid-plane has some interesting features. We observe that the density at the bilayer center for L20 and L40 are comparable to the IM density (Figs. 3A and S4) in the absence of surfactant. Further, the mid-plane density for all the three lipids is higher for laurate when compared with oleate for both compositions. The surfactant density distribution shown in Fig. S4 illustrates the influence of the difference in chain length of the two surfactants (Fig. S5). Laurate with the shorter chain length populates the central regions of both leaflets, and oleate with the longer chain length is present in the bilayer mid-plane as well.

**Fig. 3.**
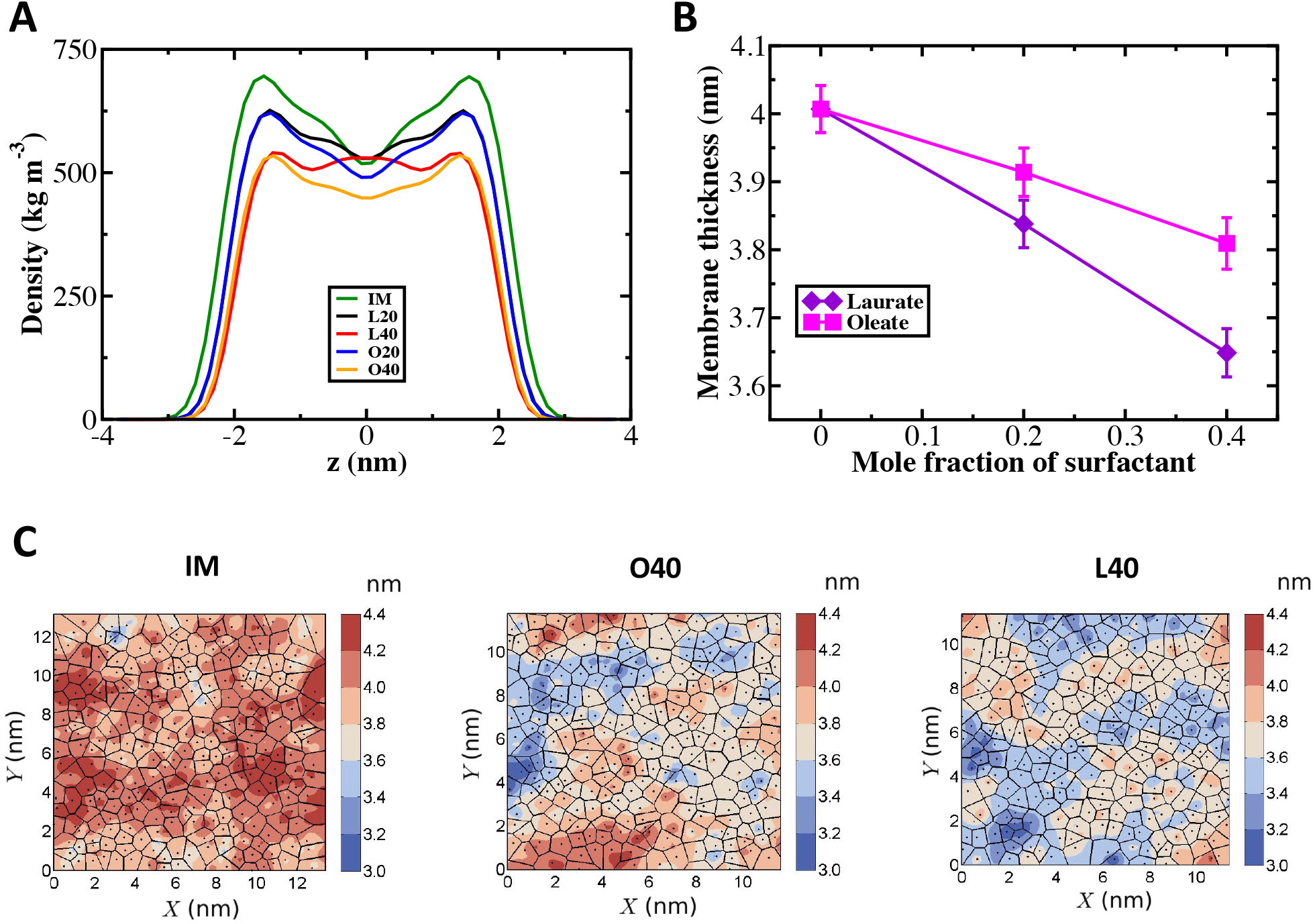
(A) Density profiles for DOPE lipids along membrane normal z-direction. (B) Membrane thickness modulation with surfactant concentration. (C) Two-dimensional Voronoi-thickness maps for IM, L40, and O40 systems.

To examine the reasons behind these trends, we computed the membrane thickness (Fig. 3B) calculated as the distance between the center of mass of the interleaflet headgroup phosphorous atoms. The bilayer thickness decreases with an increase in the surfactant concentration (Fig. 3B) for both laurate and oleate, with a greater degree of thinning observed in the case of laurate. At the highest surfactant mole fraction of 0.4, laurate induces a thinning of 8.75% when compared with a 5% reduction in the case of oleate. We have also calculated the Voronoi-thickness map^31^ for different membranes using the last configuration of the simulations (Figs. 3C and S6). We can clearly observe that bilayer thinning is more dominant in the case of laurate when compared to oleate. The trends observed in the membrane thickness suggest that the shorter chain length laurate molecules having 12 carbon atoms induce a greater extent of hydrophobic mismatch resulting in more significant membrane thinning when compared with the longer tailed oleate molecules (18 carbon atoms).

The deuterium order parameter for the lipid chains for the different surfactant concentrations is illustrated in Fig. S7. The presence of laurate results in a distinct increase in the DOPE tail disorder from C8 onward. However, in the case of oleate, the perturbation to the deuterium order parameter is far less (Fig. S7). These structural differences between the laurate and oleate mixed systems indicate that the hydrophobic mismatch induced by the incorporation of the shorter chain laurate molecule results in chain disorder and greater membrane thinning in the bacterial IM. On the contrary, oleate has an 18 carbon atom chain length which is well matched with the uniform 18 carbon atom chains of DOPE, DOPG, and TOCL lipids that make up the IM (Fig. S5).

### Electrostatics play a role in lipid-surfactant interactions

To elucidate the local structure of the membrane components in the plane of the membrane, we have calculated the lipid-lipid radial distribution function (RDF). Since the DOPE content decreases by adding surfactants, the peak heights in the RDF for the DOPE-DOPE (phosphorous atoms) interactions decreases (Fig. 4A). On the contrary, the peak heights for DOPG-DOPG interactions increases with surfactant concentration (Fig. 4B). We observed that the RDFs at a particular concentration for both laurate and oleate are similar, indicating that the different tail lengths do not perturb the correlation or relative packing between the lipid headgroups to a significant extent.

**Fig. 4.**
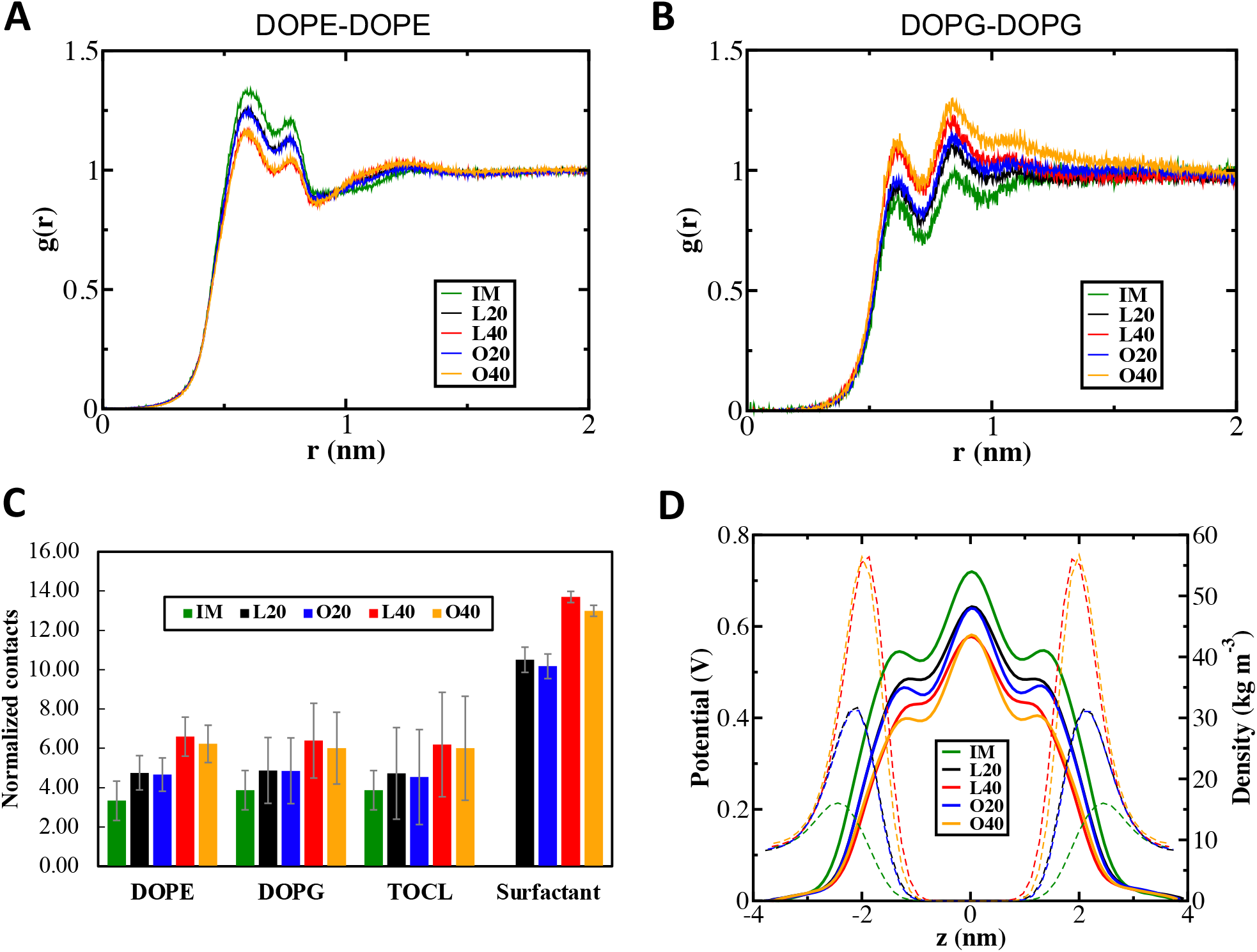
Radial distribution functions g(r) for (A) DOPE-DOPE and (B) DOPG-DOPG. (C) The normalized contacts that the headgroup of lipids (phosphorous atoms) and surfactant (oxygen atoms) make with potassium ions (details of calculation and normalization factor are provided in SI). The potassium ions within 0.4 nm of headgroups are considered for this analysis. (D) Distributions of electrostatic potential (solid lines) and the density of potassium ions (dashed lines) along membrane normal z.

To understand the trends observed in the RDF, we calculated the number of K^+^ ions within 0.4 nm cutoff of the phosphorus and oxygen head group atoms of lipids and surfactants, respectively. The number of K^+^ ions in the vicinity of the head group atoms increases as the surfactant content is increased (Fig. 4C). For the zwitterionic DOPE molecules, the presence of K^+^ due to the added surfactant does not contribute to enhanced shielding. However, in the case of the negatively charged DOPG lipids, K^+^ ions increase the electrostatic shielding between the lipids, effectively improving their relative ordering in the presence of surfactant as observed in the RDFs (Fig 4B). The trends for ion contacts are similar in DOPE, DOPG, and TOCL. The higher ion contacts with surfactant head groups, when compared with the lipids, can be attributed to both the lowered steric hindrance as well as the negative charge on the surfactant.

The electrostatic potential of the membrane decreases by incorporating anionic surfactants (Fig. 4D). The difference is dominant towards the headgroup region attributed to the charged head-groups and increased ion binding as shown by K^+^ ion density (Fig. 4D), in the presence of both surfactants. These results indicate that the change in membrane potential is expected to influence the interaction of charged species with the bacterial IMs in the presence of surfactants.

### Bending modulus and vesicle rupture

The energy associated with the local curvature of a membrane can be estimated by the bending modulus (*κ*_*c*_) using the Helfrich formulation^32^ as described in the SI. The bending modulus for the IM is estimated to be 29.1 kT, which is in good agreement with a value of 28.7 kT reported for a pure DOPE membrane^33^. This comparison is justified since the IM is mainly composed of DOPE lipids. The bending moduli for the membranes studied here reveal that the surfactant-mixed membranes are softer than the bacterial IM (Fig. 5A). A 33% decrease in *κ*_*c*_ (19.5 kT) is observed for the L20 membrane, and an extremely high reduction of 43.5% occurs for the L40 membrane with *κ*_*c*_ ∼ 16.4 kT. Oleate incorporated membranes O20 and O40 show a lower reduction in the bending modulus of 21.6% (22.8 kT) and 28.8% (20.7 kT), respectively. Hence a more significant decrease in the bending modulus can be observed in the case of laurate when compared with oleate incorporated membranes.

**Fig. 5.**
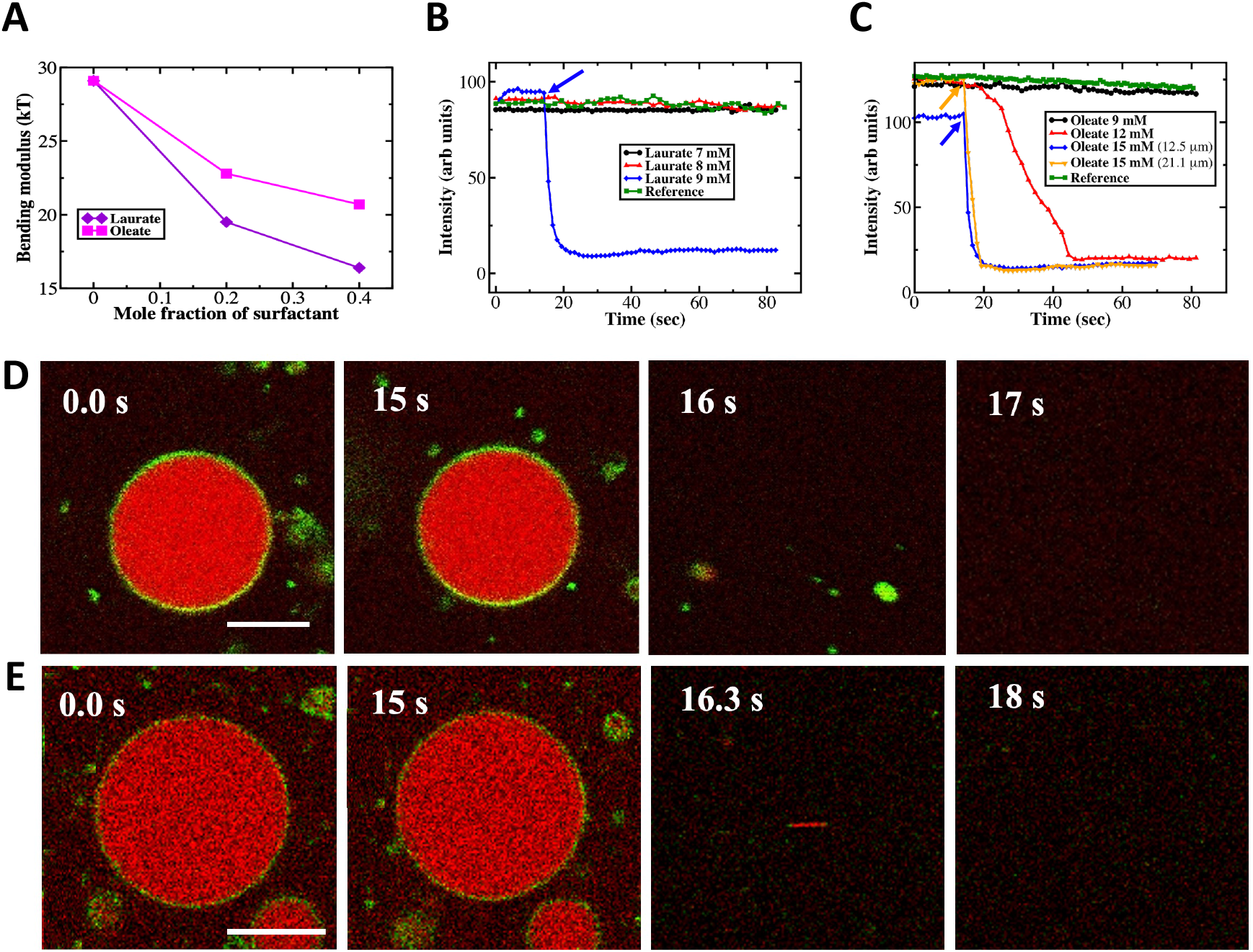
(A) Bending modulus calculated from MD simulations, for different IM systems. Fluorescence intensity versus time from GUV experiments after adding (indicated by arrows) laurate (B) and oleate (C) for the region of interest (ROI) selected, as shown in Fig. S9. Reference data is for GUVs in the absence of surfactants. Time-lapse confocal images of GUVs. (D) Addition of sodium laurate of concentration 9 mM (final concentration in the 300 *µ*l well - GUV size 7.84 *µ*m) where rapid rupture of GUVs was observed. Scale bars: 5 *µ*m. (E) Addition of sodium oleate of concentration 15 mM (final concentration in the 300 *µ*l well - GUV size 21.1 *µ*m) where rapid rupture of the GUVs occurs. Scale bars: 10 *µ*m. In the GUVs, green represents *E. coli* lipid bilayer, and the red color represents Cy5 dye filled inside the vesicles.

In order to examine the action of surfactants, we performed experiments with giant unilamellar vesicles (GUVs) composed of *E. coli* extract encapsulated with Cy5 dye. Sodium laurate was added sequentially in concentrations ranging from 2 mM to 9 mM (final concentration in the 300 *µ*l well) to the GUVS and the dye intensity from the GUVs was monitored using confocal images. In the case of sodium laurate, complete solubilization and rupture were observed at a surfactant concentration of 9 mM (Figs. 5B, D, and S9A). The observed rupture events are rapid and occur over a time scale of a few seconds (Fig. 5B). In the case of oleate, membrane solubilization and rupture were more gradual. At a surfactant concentration of 12 mM (Fig. 5C), a gradual rupture of GUVs was observed (Fig. S9B) with solubilization occurring over a period of 20 s (Fig. 5C). At 15 mM and above, rapid solubilization was observed (Figs. 5C and E). Additionally, the rupture of two independent GUVs of different sizes were found to be at similar time scales (Fig. S9). These results indicate that membrane solubilization and rupture are more effective for laurate when compared with oleate.

## Discussion

We first rationalize the differences between laurate and oleate interactions with the bacterial membranes from contact time kill (CTK) experiments performed on Gram-negative bacteria. We point out that although the experiments were carried out with the sodium salts of laurate and oleate and the MD simulations with the corresponding potassium salts, we propose a general guiding principle for the differences in their actions based on the CMC values for the different surfactants. The CTK analysis conferred better efficacy for sodium laurate at higher concentrations (> 10 mM) in agreement with previous studies on corresponding protonated fatty acids^34^, as observed by a considerable reduction in the viable population of bacteria. In contrast, the kill propensity for oleate is very weak, and we did not observe antibacterial activity up to 80 mM surfactant concentration (Fig. 1). We propose that these differences between the antimicrobial action of surfactants are driven in part by the higher CMC values (CMC^L^ = 30 mM) and shorter chain length of sodium laurate (12 carbon atoms) when compared with the lower CMC value, CMC° = 9 mM for sodium oleate. Since the free surfactant concentration (C_sf_) is limited by the CMC^35^ (C_sf_=CMC), bacteria are exposed to higher monomeric concentration of laurate when compared with oleate.

Our molecular dynamics study shows that the periplasmic PGN layer does not pose a barrier for laurate and oleate in the monomeric state. However, the number of events for translocation of a single laurate molecule across the PGN layer was much higher, indicating faster kinetics of translocation in the case of laurate when compared to oleate. We report that these differences are based on the interplay between interactions of surfactant molecules with sugar and amino acids subunits of the PGN layer. Additionally, the PGN layer offers a barrier to the aggregates of both surfactants, though the tendency for aggregate formation causes differences in the potency. The higher CMC value of sodium laurate (30 mM) results in an increased single-molecule population^22^ above the CMC. Further, if the concentration regime below the CMC is sufficient to induce antimicrobial activity, the greater bacterial kill will occur, as was observed for laurate in the CTK data. In the case of sodium oleate having a much lower CMC value (CMC° = 9 mM), micelles formed at low concentrations are unable to translocate through the PGN layer, as illustrated in our MD simulations. For the case of oleate, if the antimicrobial activity sets in above the CMC of 9 mM, the formation of micelles prevents the concentration of free surfactant from exceeding the CMC value. The differences in the barrier offered by the PGN layer for these molecules at higher concentrations explain the differences in the efficacy of these molecules against *E. coli* (Fig. 6).

**Fig. 6.**
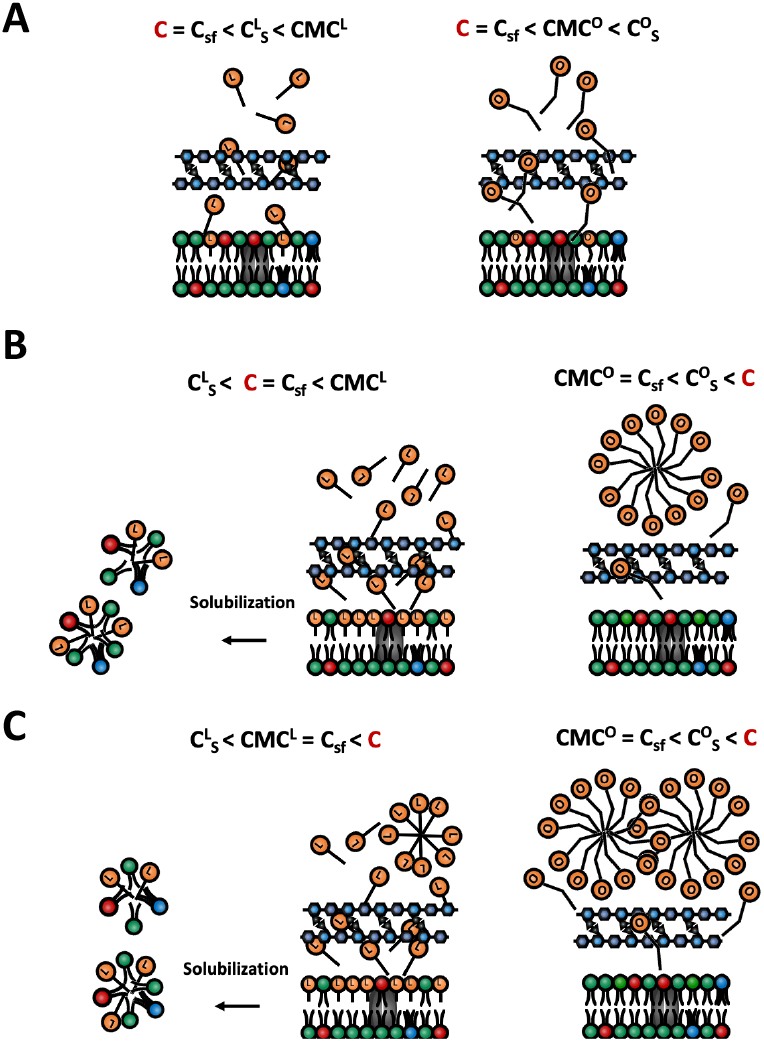
Schematic illustration of the translocation and solubilization mechanism for the surfactant molecules based on their concentrations and CMC values. C and C_sf_ are the total and free surfactant concentrations, respectively. Critical micelle concentrations for laurate and oleate are represented by CMC^L^ and CMC°, respectively. Laurate and oleate concentration required for inner membrane solubilization are represented by C^L^_S_ and C°_S_, respectively. (A) Low surfactant concentration with no killing activity, (B) Intermediate concentration where single laurate molecules are active, and (C) both laurate and oleate form micelles, however only laurate is active due to sufficient single molecule surfactant concentration.

In addition to the differences in the mechanism and barriers offered to both the surfactants by the PGN layer, the IM interactions also play a crucial role. Our surfactant incorporated IM atomistic models were found to be stable over the course of the simulation and we did not observed any ripple formation as observed in previous studies of surfactant/co-surfactant membranes^36^. However, among the key observations, the lipid tails become increasingly disordered, and the bilayers show a reduction in the membrane thickness with rising surfactant concentration. In general, these changes were greater and more accentuated in the case of the shorter chain laurate molecules when compared with oleate. We attributed the observed differences between laurate and oleate primarily to the increased hydrophobic mismatch for laurate which is 12-carbon long, while the lipids chains are 18-carbons in length, similar to that of oleate. The presence of laurate decreases the correlations between DOPE molecules to a greater extent when compared with oleate, indicating the greater disorder induced due to the increased hydrophobic mismatch. An opposite effect in the correlations is observed for DOPG molecules which are smaller in content in the IM. The changes in the charge interactions are attributed to higher ion binding with increasing surfactant concentrations. This decreases the membrane potential and can influence the interaction of charged species like cationic antimicrobial peptides with the IM. This could also explain the reason for synergistic effects exhibited by the surfactants with cationic peptides^37^. The change in membrane potential can also affect the local organization of proteins involved in cell division^38^. Laurate is also found to decrease the bending modulus of the IM to a greater extent when compared with oleate. Our MD simulations indicate that laurate is able to induce a greater extent of disruption to the IM when compared with oleate, consistent with increased antibacterial activity observed for laurate in both the CTK data as well as the GUV rupture data.

The GUV experiments provided a closer correspondence to interpret findings from the IM molecular dynamics simulations. Oleate caused GUV rupture at higher concentrations (> 12 mM) when compared with laurate (> 9 mM). Based on GUV experiments, it can be inferred that the minimum surfactant concentration required for IM solubilization is lower in the case of laurate (C^L^_S_ = 9 mM) as compared to oleate (C°_S_ = 12 mM). Note that, C^L^_S_ = 9 mM is much lower than the corresponding CMC value of sodium laurate (30 mM), however C°_S_ = 12 mM is only slightly larger than the CMC (9 mM) for sodium oleate. These results clearly show the increased tendency for laurate to induce membrane solubilization and rupture when compared with the longer chain oleate.

Our study provides compelling evidence for the differences in surfactant action on the IM, and MD simulations illustrate the interactions of both laurate and oleate with the PGN layer. However, the interaction of the surfactants with the OM is unclear, and here we discuss the possibilities of membrane solubilization and transport through OM channels. Surfactants can solubilize bacterial membranes, and several mechanisms have been discussed in the literature^4^. If laurate action is driven by solubilization of the OM, then it is likely that oleate would also follow a similar mode of action. Since the concentration of free surfactant is limited by its CMC, the lower monomeric concentration for oleate could preclude the OM solubilization if this indeed was the primary mechanism of antibacterial action. Additional investigations would have to be carried out to determine the interactions and possible solubilization action of surfactants on the bacterial OM. We cannot rule out the possibility that antibacterial activity can be driven by a combination of the outer membrane and inner membrane solubilization, and given the complex architecture of the outer membrane with the increased interaction between long chain lipopolysaccharides, the solubilization kinetics is expected to be slow when compared to that of the IM. In either event, the availability of surfactant to the PGN and IM would be dependent on the CMC.

Surfactants could access the IM through defects created in the OM, or one could also speculate transport of surfactants across the outer membrane through the OM transporters. In this regard, the FadL transporters^39^ are known to allow the transport of fatty acids. Surfactants can access the OM of bacteria through these transporters and reach the periplasmic space where the PGN layer is located. Since both oleate and laurate molecules are able to pass through the PGN layer, albeit at different rates, they would be able to access the IM. Solubilization would then follow, provided the concentration of surfactant at the IM is sufficiently high. This hypothesis which supports the view that surfactant action occurs at the IM is also corroborated by the good agreement between the C^L^_S_ (9 mM) observed in GUV rupture and the threshold concentration (10 mM) for observing a one log reduction in viable bacterial population in the case of sodium laurate. This supports the notion that laurate molecule concentration at the IM is sufficiently high to cause solubilization of the IM and kill the bacteria. With regard to oleate, we argue that if oleate molecules access the IM, its concentration is limited by the CMC of 9 mM, which is below the threshold concentration to induce solubilization and rupture the IM. This fact is also supported by the higher concentration required by oleate to solubilize GUVs.

We summarize our findings in Fig. 6 which provides a schematic illustration of the aforementioned mechanism, which assumes that surfactants are able to access the IM and induce membrane rupture and damage. If the free surfactant concentration, C_sf_ is below the respective CMC values as well as below the IM solubilization concentrations for laurate (C^L^_S_) and oleate (C°_S_), kill activity is absent (Fig. 6A). Additionally, if C_sf_ is above solubilization concentration and below CMC, then kill activity would occur as illustrated in Fig. 6B. The aforementioned kill mechanism is expected for surfactants like laurate. While in the case of surfactants like oleate where, C°_S_ > CMC°, C_sf_ will not exceed CMC° hence limiting its kill activity (Figs. 6B and C). Laurate however will be effective even above CMC^L^, as the C_sf_ is greater than the corresponding C^L^_S_.

## Conclusions

Using a combination of experiments and molecular dynamics simulations, we differentiate between the action of surfactants, whose primary differences lie in the hydrocarbon tail lengths, with both the IM and intervening PGN layer of Gram-negative bacteria. Our study reveals that the peptidoglycan cell wall, generally perceived to be a passive barrier for transport, serves as a filter modulating the translocation of surfactants based on their physicochemical and aggregation properties, providing insights into a hitherto unexplored regime of transport across the bacterial cell wall. We specifically provide insights into the modulation of the IM structural and mechanical properties as a function of surfactant chemistry - in this case laurate versus oleate. The critical concentration required for bacterial kill is related to the CMC, which determines the availability of surfactant at the membrane interface. Additional investigations would be required to confirm the role of protein channels for surfactant transport as well as the tendency for surfactants to solubilize the OM of Gram-negative bacteria. Our study provides a quantitative framework to assess the antibacterial efficacy of surfactant molecules. The methods and insights presented here can be extended to evaluate interactions of other surface active molecules and optimize antibacterial formulations.

## Author contributions

Conceptualization: P.S. and K.G.A; Data curation: P.S.; Funding acquisition: J.S.R, J.K.B and K.G.A.; Project administration: P.S. and K.G.A.; Resources: M.W., J.S.R, J.K.B and K.G.A; MD Simulations: P.S. and R.K.V.; GUV experiments: S.P.; Contact time kill assays: N.P.; Supervision: M.W., J.S.R, J.K.B and K.G.A; Writing - original draft: P.S., R.K.V., S.P. and N.P.; Writing - review & editing: P.S., R.K.V., J.S.R., J.K.B and K.G.A.

## Acknowledgements

We thank Unilever Research and Development (Bangalore, India) for funding this research and Scott Singleton for valuable discussions. We also thank the Supercomputer Education and Research Center (SERC) for availing computational facility at the Indian Institute of Science, Bangalore, and the Department of Science and Technology, India for funding computational resources used for parts of this study.

## Notes and references

1 N. A. Falk, Journal of Surfactants and Detergents, 2019, 22, 1119–1127.

2 B. C. Hoefler, K. V. Gorzelnik, J. Y. Yang, N. Hendricks, P. C. Dorrestein and P. D. Straight, Proceedings of the National Academy of Sciences, 2012, 109, 13082–13087.

3 D. K. F. Santos, R. D. Rufino, J. M. Luna, V. A. Santos and L. A. Sarubbo, International journal of molecular sciences, 2016, 17, 401.

4 H. Heerklotz, Quarterly reviews of biophysics, 2008, 41, 205.

5 W. Im and S. Khalid, Annual Review of Physical Chemistry, 2020, 71, 171–188.

6 S. Zhang, S. Ding, J. Yu, X. Chen, Q. Lei and W. Fang, Langmuir, 2015, 31, 12161–12169.

7 B. K. Yoon, J. A. Jackman, E. R. Valle-González and N.-J. Cho, International journal of molecular sciences, 2018, 19, 1114.

8 T. Nakatsuji, M. C. Kao, J.-Y. Fang, C. C. Zouboulis, L. Zhang, R. L. Gallo and C.-M. Huang, Journal of Investigative Dermatology, 2009, 129, 2480–2488.

9 C. B. Huang, Y. Alimova, T. M. Myers and J. L. Ebersole, Archives of oral biology, 2011, 56, 650–654.

10 A. Laatiris, M. El Achouri, M. R. Infante and Y. Bensouda, Microbiological Research, 2008, 163, 645–650.

11 B. H. Morrow, P. H. Koenig and J. K. Shen, Langmuir, 2013, 29, 14823–14830.

12 J. J. Janke, W. D. Bennett and D. P. Tieleman, Langmuir, 2014, 30, 10661–10667.

13 X. Zhang, F. Song, M. Taxipalati, W. Wei and F. Feng, PLoS One, 2014, 9, e114845.

14 N. Joondan, S. Jhaumeer-Laulloo, P. Caumul and M. Akerman, Journal of Physical Organic Chemistry, 2017, 30, e3675.

15 P. Sharma, S. Parthasarathi, N. Patil, M. Waskar, J. S. Raut, M. Puranik, K. G. Ayappa and J. K. Basu, Langmuir, 2020, 36, 8800–8814.

16 J. Michel, Y. Wang, I. Kiesel, Y. Gerelli and V. Rosilio, Langmuir, 2017, 33, 11028–11039.

17 N. Paracini, L. A. Clifton and J. H. Lakey, Biochemical Society Transactions, 2020, 48, 2139–2149.

18 N. K. Sarangi, K. Ayappa, S. S. Visweswariah and J. K. Basu, Langmuir, 2016, 32, 9649–9657.

19 T. S. Carpenter, J. Parkin and S. Khalid, The journal of physical chemistry letters, 2016, 7, 3446–3451.

20 R. Vaiwala, P. Sharma, M. Puranik and K. G. Ayappa, Journal of Chemical Theory and Computation, 2020, 16, 5369–5384.

21 W. Shinoda, R. DeVane and M. L. Klein, Soft Matter, 2008, 4, 2454–2462.

22 D. T. King, D. B. Warren, C. W. Pouton and D. K. Chalmers, Langmuir, 2011, 27, 11381–11393.

23 S. Bandyopadhyay, J. C. Shelley and M. L. Klein, The Journal of Physical Chemistry B, 2001, 105, 5979–5986.

24 A. Pizzirusso, A. De Nicola, G. A. Sevink, A. Correa, M. Cascella, T. Kawakatsu, M. Rocco, Y. Zhao, M. Celino and G. Milano, Physical Chemistry Chemical Physics, 2017, 19, 29780–29794.

25 M. J. Abraham, T. Murtola, R. Schulz, S. Páll, J. C. Smith, B. Hess and E. Lindahl, SoftwareX, 2015, 1, 19–25.

26 J. C. Gumbart, M. Beeby, G. J. Jensen and B. Roux, PLoS Comput Biol, 2014, 10, e1003475.

27 S. Jo, T. Kim, V. G. Iyer and W. Im, Journal of computational chemistry, 2008, 29, 1859–1865.

28 J. B. Klauda, R. M. Venable, J. A. Freites, J. W. O’Connor, D. J. Tobias, C. Mondragon-Ramirez, I. Vorobyov, A. D. MacKerell Jr and R. W. Pastor, The journal of physical chemistry B, 2010, 114, 7830–7843.

29 S. R. Durell, B. R. Brooks and A. Ben-Naim, The Journal of Physical Chemistry, 1994, 98, 2198–2202.

30 I. I. Ponmalar, R. Cheerla, K. G. Ayappa and J. K. Basu, Proceedings of the National Academy of Sciences, 2019, 116, 12839–12844.

31 P. Sharma, R. Desikan and K. G. Ayappa, bioRxiv, 2020.

32 W. Helfrich, Zeitschrift für Naturforschung A, 1978, 33, 305–315.

33 R. M. Venable, F. L. Brown and R. W. Pastor, Chemistry and physics of lipids, 2015, 192, 60–74.

34 C. P. Churchward, R. G. Alany and L. A. Snyder, Critical reviews in microbiology, 2018, 44, 561–570.

35 K.-C. Huang, C.-M. Lin, H.-K. Tsao and Y.-J. Sheng, The Journal of chemical physics, 2009, 130, 06B622.

36 A. Debnath, F. M. Thakkar, P. K. Maiti, V. Kumaran and K. Ayappa, Soft Matter, 2014, 10, 7630–7637.

37 K. Liu, L. Yang, X. Peng, H. Gong, J. Wang, J. R. Lu and H. Xu, Langmuir, 2020, 36, 3531–3539.

38 H. Strahl and L. W. Hamoen, Proceedings of the National Academy of Sciences, 2010, 107, 12281–12286.

39 M. C. Wiener and P. S. Horanyi, Proceedings of the National Academy of Sciences, 2011, 108, 10929–10930.

