## Supplementary Information for "Interactions of surfactants with the bacterial cell wall and inner membrane: Revealing the link between aggregation and antimicrobial activity"

**This PDF file includes:**

Supplementary text
Figs. S1 to S9
Tables S1 to S2
SI References

S2 The evolution of the number of clusters in simulations carried out with the peptidoglycan layer. L10 and L30 contain 10 and 30 potassium laurate molecules respectively and O10 and 030 correspond to the oleate system. A single surfactant is counted as one cluster. Once a molecule is within 0.35 nm of another molecule or aggregate, it is counted as part of the aggregate. The analysis indicates that the L10 system does not form clusters and we see formation of dimers and trimers, however the L30 system continuously evolves toward a large aggregate. At the end of the 500 ns simulation we have a monomer and a large aggregate. In the case of the oleate systems aggregate formation is rapid and a single aggregate is observed for O10 while two aggregates were observed for the O30 system. . . . 8

S3 Simulation snapshots showing bacterial inner membrane systems IM, L20, L40, O20 and O40. Lipid phosphorus and surfactant oxygen atoms are represented by van der Waals spheres in pink and purple, respectively. Phospholipids DOPE (green), DOPG (red), cardiolipin (blue) and surfactants (orange) are represented by bonds. 9

S4 Density distributions of lipids and surfactants. The increased density of lipids at the bilayer mid-plane in the case of L20 and L40 indicates the presence of interdigitation. The density of laurate in L20 and L40 is negligible at the mid plane of bilayer, which can be attributed to the shorter length of the surfactant with respect to the lipids resulting in a greater hydrophobic mismatch. . . . 10

S5 Molecular structures for laurate, oleate, DOPE, DOPG and TOCL1. . . . 11

S6 Two-dimensional Voronoi-thickness maps, thickness is measured as a distance between headgroup phosphorus atoms across the leaflets. Thinning can be observed in the membranes and this effect is greater in the laurate systems, L20 and L40 when compared with the oleate systems, O20 and O40. . . . 12

S7 Deuterium order parameter, $S_{cd}$ for the DOPE SN1, DOPE SN2, DOPG SN1, DOPG SN2, TOCL1 A, TOCL1 B, TOCL1 C, TOCL1 D and surfactant chains. The carbon atom numbers (x-axis) are counted from the carbon atom labelled as shown in Figure S5 for the different molecules. In all cases we find greater disorder induced for the laurate (L20 and L40) systems when compared with the corresponding oleate systems (O20 and O40). Additionally the disorder is greatest toward the membrane mid-plane. In the case of the surfactants greater disorder is observed for the laurate molecules due to the increased hydrophobic mismatch. . . . . . . . . . . . 13

S8 Static structure factor for membrane height fluctuations. The corresponding values of bending modulus are given in main text. Our simulation box sizes are sufficiently large to capture the low $q$ regime in order to extract the bending modulus. . . . . . . . . . . . . . . . . . . . . . . . . . . . . . . . . . . . . . . . . . . . . . 14

S9 Time-lapse images of multiple GUVs. (A) Addition of sodium laurate of concentration 9 mM (final concentration in the 300 $\mu$l well - GUV size 7.84 $\mu$m) - rupture of GUVs was observed. (B) Addition of sodium oleate of concentration 12 mM (final concentration in the 300 $\mu$l well - GUV size 10.7 $\mu$m) - makes the GUVs to get slowly ruptured. (C) Addition of sodium oleate of concentration 15 mM (final concentration in the 300 $\mu$l well - GUV sizes 12.5 $\mu$m and 21.1 $\mu$m) - makes the GUVs to get ruptured immediately. . . . . . . . . . . . . . . . . . 15

### Supporting Information Text

### SI methods

**Prepartation and fixing of GUVs.** *E. coli* total lipid extract (hereafter referred to as *E. coli* lipids) was purchased from Avanti Polar Lipids and used without further purification. Fluorescent markers ATTO 488 1,2-Bis(dimethylphosphino)ethane (DMPE), head tagged lipid (ATTO-Tec GmbH) and Cy5-NHS Ester dissolved in tris buffer used to fill the GUVs were purchased from Sigma-Aldrich and Thermo Fischer scientific respectively. *E. coli* inner membrane (IM) GUVs were prepared by the electroformation method (1, 2). Briefly, 2 mM of *E. coli* lipid with Atto 488 DMPE and Biotinylated-DMPE of required ratio, were spread uniformly on a cleaned ITO coated glass slide. The lipids were dried by desiccating the slides in a vacuum desiccator for 2 hours. 1.5 ml of deionized (DI) water with 250 mM sucrose was added with 200 nM Cy5 and added between the ITO slides attached with the polydimethylsiloxane (PDMS) spacer of 2 mm thickness. ITO slides were connected to the frequency generator with a sine wave at the frequency of 10 Hz, output voltage of 1 Vpp with no phase difference. The slides were placed inside the incubator at a 55°C for 90 min. Using a 1 ml pipette, the solution was carefully mixed and stored in the micro-centrifuge tubes at 4°C until imaging. This procedure yielded good number of GUVs. In order to fix the GUVs on the glass substrate, DPPC bilayer was prepared by vesicle fusion method with Biotinyl PE in the ratio 500:1. Small unilamellar vesicles (SUV) of DPPC of 1 mM was prepared by sonication method in 1X PBS buffer. In order to prepare the bilayer, desired amount of SUVs were added to plasma treated coverslip. It was incubated at 55 ° C for 1 hour and then the slide was washed with water thrice to remove unfused vesicles. Streptavidin was added to the slide and left for 30 min incubation at room temperature. The unbound streptavidin was washed with DI water. Then the GUVs were added to the bilayer and diluted with 250 mM glucose buffer during imaging. In this method many GUVs were attached to the DPPC bilayer through strong covalent interactions between streptavidin-biotinylated PE. The sample was gently washed later with buffer to remove the free suspended vesicles.

**Table S1. Details of MD Simulations performed to observe interactions of surfactants with peptidoglycan layer**

| System | Laurate | Oleate | Simulation time (ns) | Size ($L_x$ × $L_y$ × $L_z$ $nm^3$) | Total atoms | TIP3P | K$^+$ ions |
|---|---|---|---|---|---|---|---|
| L1 | 1 | 0 | 500 | 13.71 × 13.71 × 7.20 | 138,358 | 42,158 | 183 |
| L10 | 10 | 0 | 500 | 13.71 × 13.71 × 7.17 | 138,187 | 41,987 | 192 |
| L30 | 30 | 0 | 500 | 13.71 × 13.71 × 7.09 | 136,421 | 41,145 | 212 |
| O1 | 0 | 1 | 500 | 13.71 × 13.71 × 7.20 | 138,305 | 42,135 | 183 |
| O10 | 0 | 10 | 500 | 13.71 × 13.71 × 7.16 | 137,861 | 41,825 | 192 |
| O30 | 0 | 30 | 500 | 13.71 × 13.71 × 6.90 | 132,365 | 39,633 | 212 |

**Molecular dynamics simulations.** The systems were energy minimized and equilibrated for 10 ns and 50 ns for peptidoglycan (PGN) and lipid membranes, respectively, by following standard CHARMM-GUI protocols. In all cases, equilibration was achieved by monitoring the lateral membrane area over the course of the simulation. The periodic boundary conditions are used in all three directions. Atomic velocities and positions were updated with a time step of 2 fs employing leapfrog integrator. All simulations were performed at 303.15 K using a Nosé-Hoover thermostat (3) with a coupling constant of 1 ps. Semi-isotropic pressure coupling was employed using Parrinello-Rahman barostat (4) with a coupling constant of 5 ps for lipid membrane systems. The particle-mesh Ewald (PME) (5) was chosen to calculate long-range electrostatic interactions with a cutoff of 1.2 nm, while the linear constraint solver (LINCS) (6) was used for constraining hydrogen bonds. Verlet cutoff scheme is chosen to compute the non bonded interactions (Lennard-Jones 6-12), which are truncated at the cutoff radius of 1.2 nm, and shifted with a force-switch function between 1.0 to 1.2 nm.

**Table S2. Compositions of lipid membranes used in MD Simulations**

| System | Total lipids | PE | PG | CL | Laurate | Oleate | Simulation time(ns) | Size ($L_x \times L_y \times L_z\ nm^3$) | Total atoms | TIP3P | $K^+$ ions |
|---|---|---|---|---|---|---|---|---|---|---|---|
| IM | 512 | 384 | 104 | 24 | 0 | 0 | 1000 | 12.95 × 12.95 × 7.39 | 127,515 | 19,417 | 128 |
| L20 | 408 | 306 | 82 | 20 | 104 | 0 | 1000 | 12.35 × 12.35 × 7.71 | 120,840 | 20,530 | 206 |
| L40 | 308 | 232 | 60 | 16 | 204 | 0 | 1000 | 11.64 × 11.64 × 7.47 | 104,092 | 18,164 | 280 |
| O20 | 408 | 306 | 82 | 20 | 0 | 104 | 1000 | 12.50 × 12.50 × 7.61 | 122,420 | 20,502 | 206 |
| O40 | 308 | 232 | 60 | 16 | 0 | 204 | 1000 | 11.87 × 11.87 × 7.40 | 107,104 | 18,080 | 280 |

**Analysis and Visualization.** Visual molecular dynamics 1.9.3 (VMD) (7) is used for rendering MD snapshots. Membrane thickness is computed using the MEMBPLUGIN 1.1 tool (8) in VMD. The Voronoi-thickness modulation maps are plotted using the 'Voronoi' function available in MATLAB (https://www.mathworks.com) for the x, y coordinates (upper leaflet) of lipid phosphorus atom from the corresponding MD snapshot. Cluster analysis, density distribution, radial distribution function, membrane average thickness, normalized contacts, deuterium order parameter, and membrane electrostatic potential were computed using GROMACS (9) post processing tools.

**Algorithm for calculating translocation.** We have defined three regions along the z-direction in our simulation box using the center of mass of the PGN layer as a reference. The molecule trajectory along the z-direction (parallel to the membrane normal) has been flagged as U (upper), L (lower), and PG (buffer) zone with respect to the PGN layer (Fig. S1). The buffer zone is a 2 nm thick region around the PGN layer center of mass to ensure that molecule interactions are negligible with the PGN layer once it crosses this region. A translocation event is determined as the passage of the molecule from the upper to the lower zone or vice versa, through the buffer area and not through periodic boundaries. Fig. S1 shows a few cases where green and blue trajectories represents translocation and no translocation events, respectively.

**Normalized lipid-ion contacts.** We have calculated the number of $K^+$ ions within 0.4 nm cutoff of the phosphorus and oxygen head group atoms of lipids and surfactants, respectively. The values obtained from this analysis are normalized with respect to the number of molecules and the number of ions in the system.

$$\text{Normalized contacts} = \frac{10000 \times \text{Total number of contacts}}{\text{N}_F \times \text{N}_M \times \text{N}_K} \qquad [1]$$

where $\text{N}_F$ is the number of frames considered for this analysis, $\text{N}_M$ is the total number of lipids or surfactants, and $\text{N}_K$ is the total number of potassium ions.

**Membrane electrostatic potential.** The electrostatic potential (V) of the membrane, can be calculated by carrying out the double integral of charge density $\rho(z)$, using the following expression:

$$V(z) = -\frac{1}{\varepsilon_0}\int_0^z dz' \int_0^{z'} \rho(z'')\, dz'' \quad [2]$$

where $\varepsilon_0$ is the free space permittivity, and $z = 0$ represent the membrane mid plane.

**Bending modulus calculation.** Bending modulus ($\kappa$), which is a mechanical property of the membrane was calculated using Helfrich(10) formulation, the static structure factor corresponding to the thermal fluctuations of the local membrane-height scales with low wave modes $q$ as $q^{-4}$ according to,

$$S(q) = k_B T/\kappa q^4 \quad [3]$$

Here $k_B$ denotes the Boltzmann constant, while T is the absolute temperature. With a membrane area of size $A$, the bending modes $q = \frac{2}{\pi\sqrt{A}}(n_x, n_y)$, where $n_x$ and $n_y$ are the integer numbers. Figure S8 shows the low bending modes following the $q^{-4}$ scaling. The bending modulus is deduced by fitting the structure factor data to equation 3.

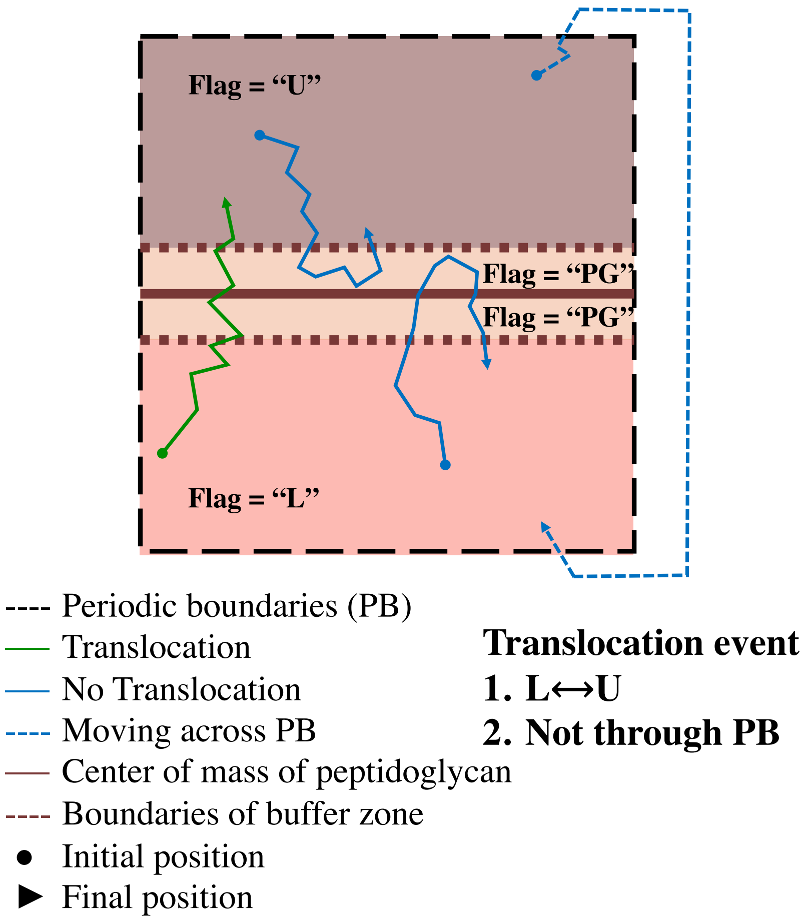

**Fig. S1.** Schematic showing the algorithm to count number of translocation events which occur when a surfactant molecule crosses the PGN layer as illustrated. The green and blue trajectories represent translocation and no translocation events, respectively.

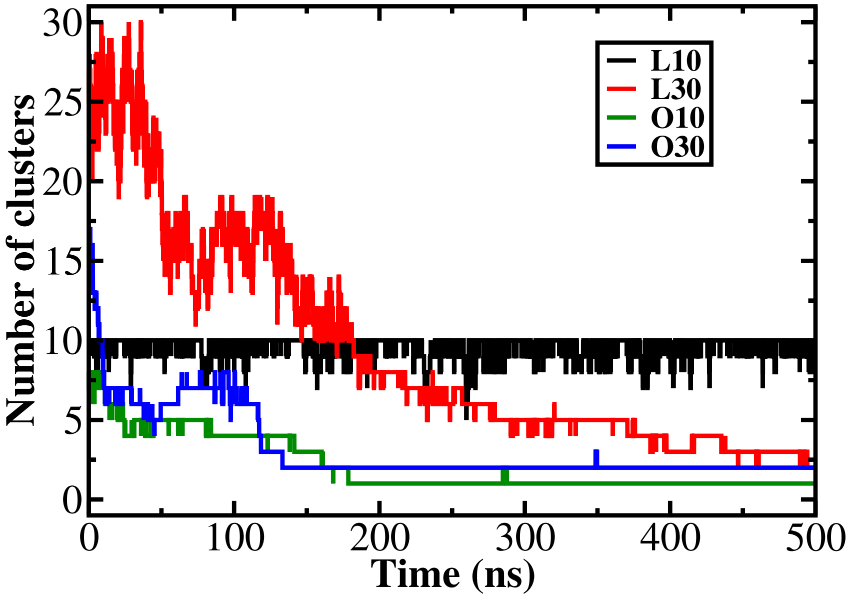

**Fig. S2.** The evolution of the number of clusters in simulations carried out with the peptidoglycan layer. L10 and L30 contain 10 and 30 potassium laurate molecules respectively and O10 and 030 correspond to the oleate system. A single surfactant is counted as one cluster. Once a molecule is within 0.35 nm of another molecule or aggregate, it is counted as part of the aggregate. The analysis indicates that the L10 system does not form clusters and we see formation of dimers and trimers, however the L30 system continuously evolves toward a large aggregate. At the end of the 500 ns simulation we have a monomer and a large aggregate. In the case of the oleate systems aggregate formation is rapid and a single aggregate is observed for O10 while two aggregates were observed for the O30 system.

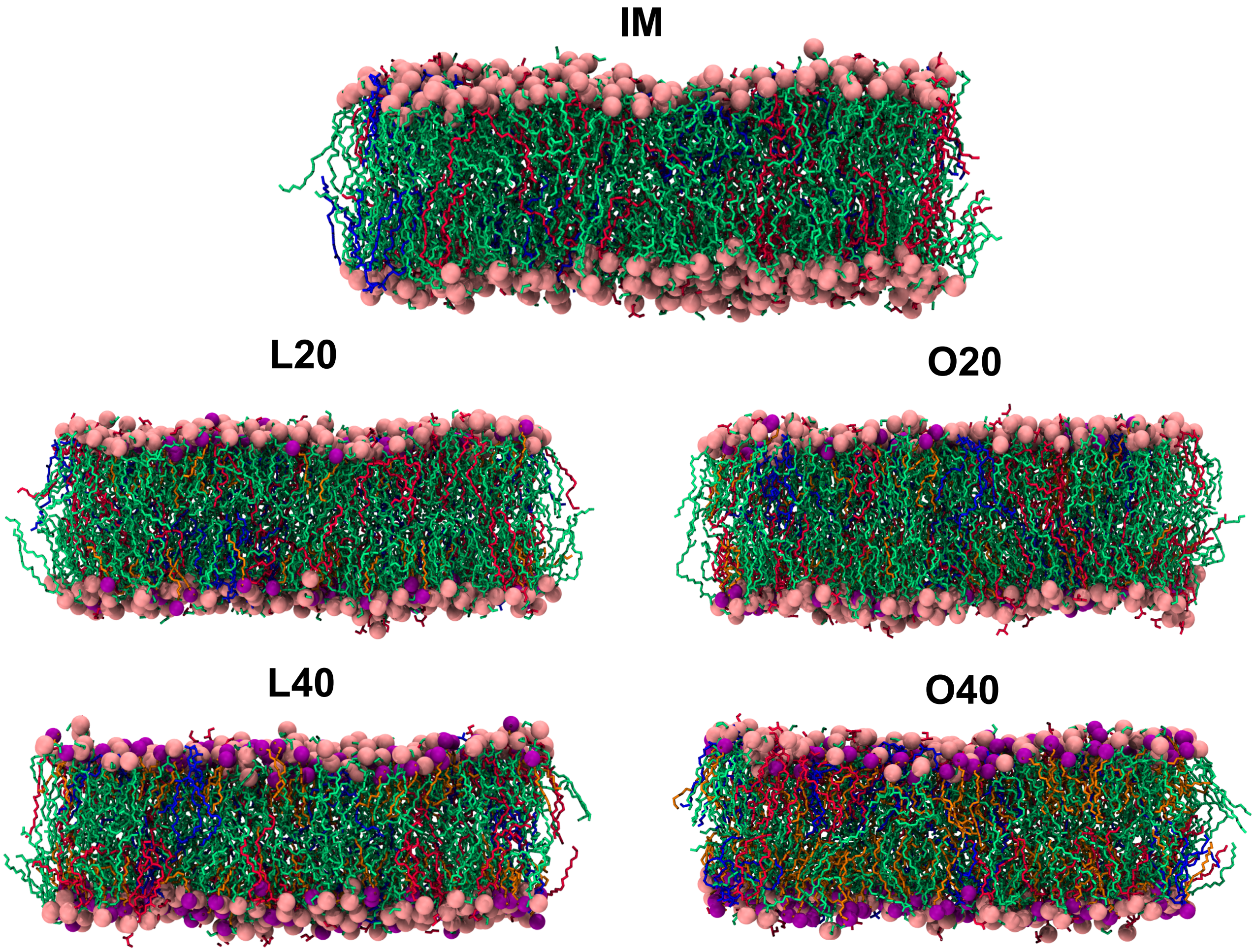

**Fig. S3.** Simulation snapshots showing bacterial inner membrane systems IM, L20, L40, O20 and O40. Lipid phosphorus and surfactant oxygen atoms are represented by van der Waals spheres in pink and purple, respectively. Phospholipids DOPE (green), DOPG (red), cardiolipin (blue) and surfactants (orange) are represented by bonds.

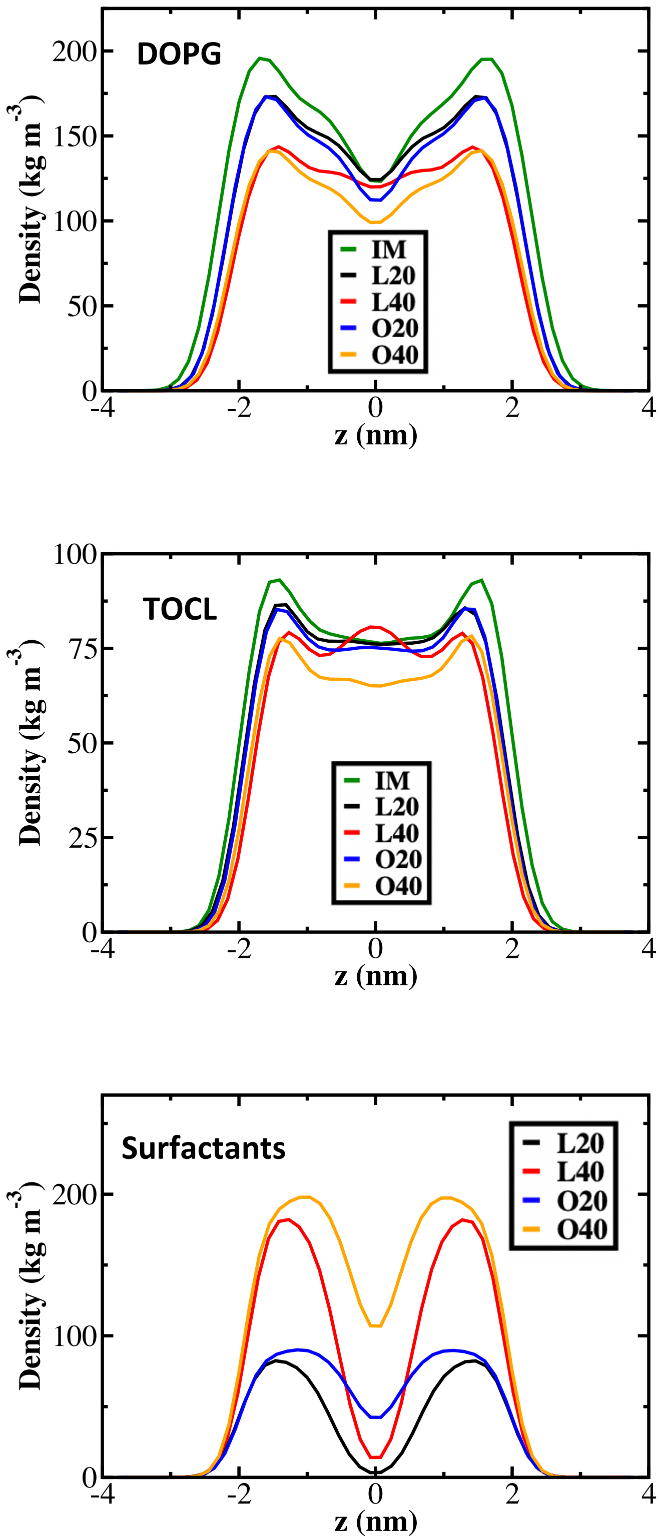

**Fig. S4.** Density distributions of lipids and surfactants. The increased density of lipids at the bilayer mid-plane in the case of L20 and L40 indicates the presence of interdigitation. The density of laurate in L20 and L40 is negligible at the mid plane of bilayer, which can be attributed to the shorter length of the surfactant with respect to the lipids resulting in a greater hydrophobic mismatch.

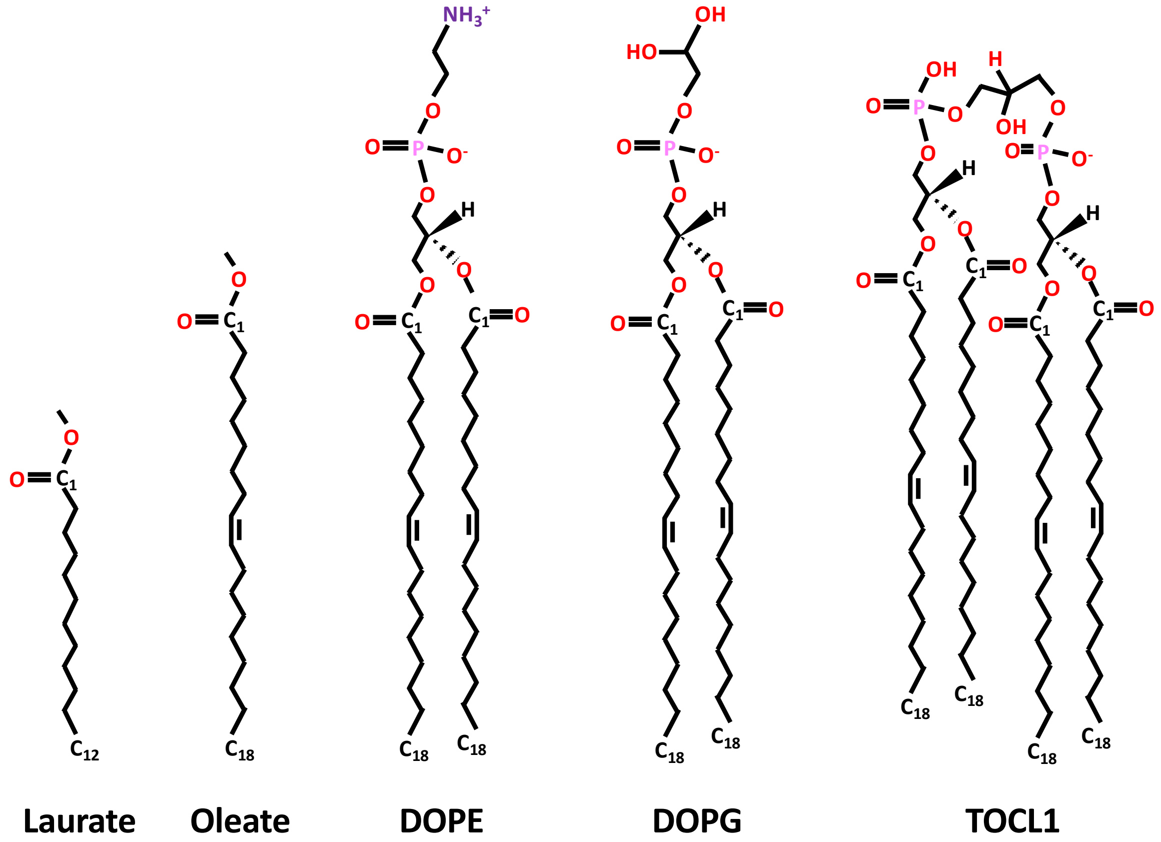

**Fig. S5.** Molecular structures for laurate, oleate, DOPE, DOPG and TOCL1.

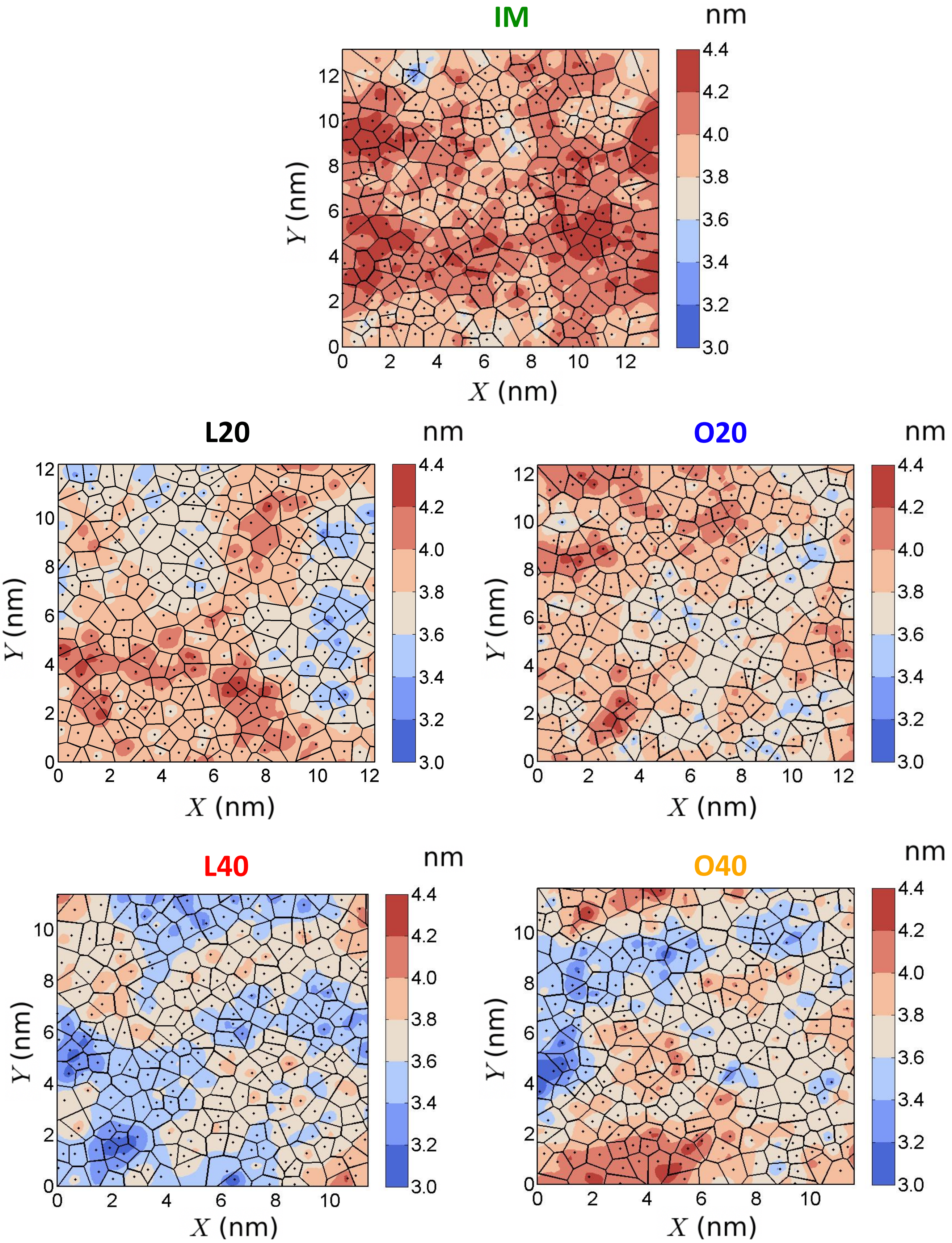

**Fig. S6.** Two-dimensional Voronoi-thickness maps, thickness is measured as a distance between headgroup phosphorus atoms across the leaflets. Thinning can be observed in the membranes and this effect is greater in the laurate systems, L20 and L40 when compared with the oleate systems, O20 and O40.

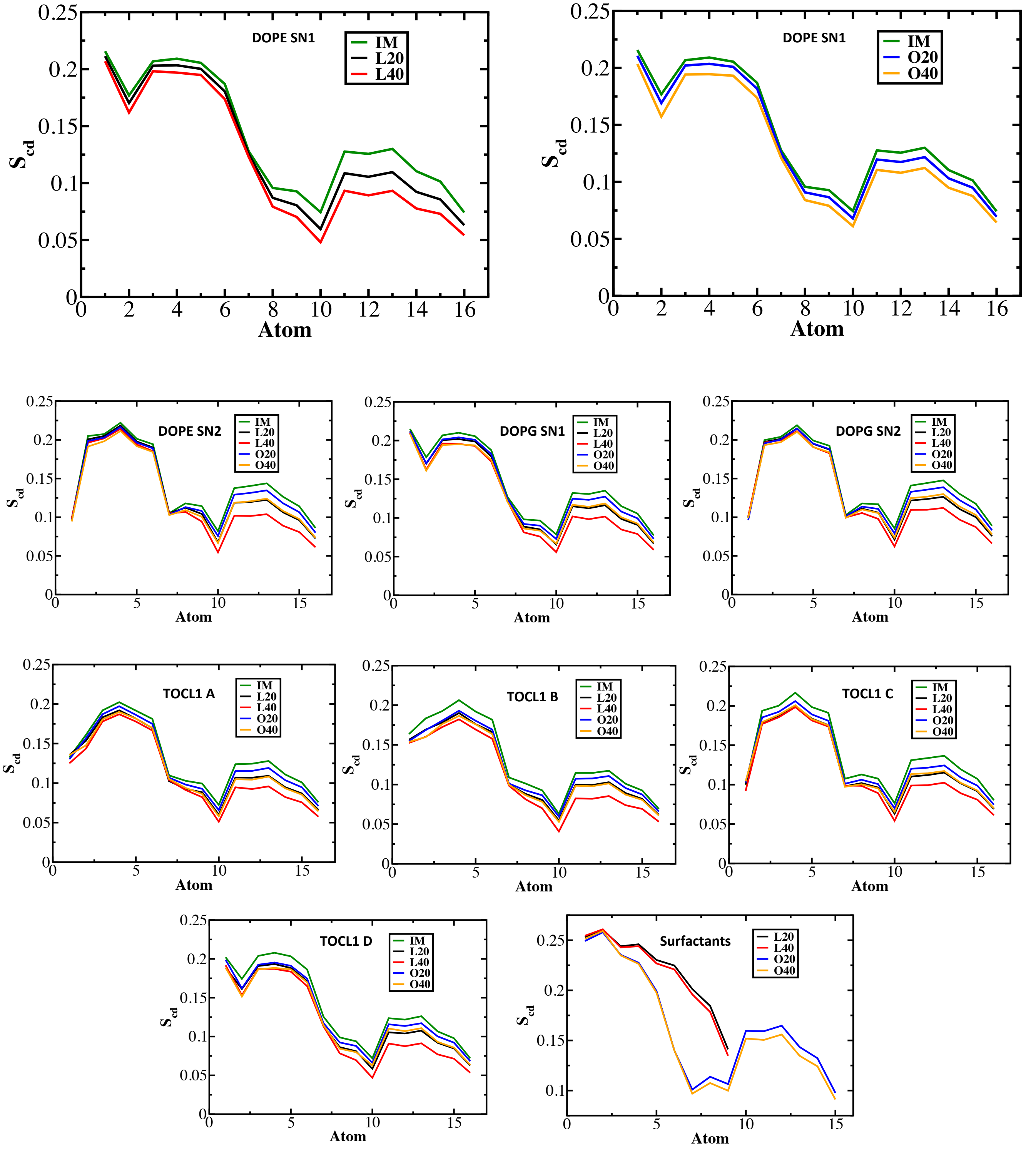

**Fig. S7.** Deuterium order parameter, $S_{cd}$ for the DOPE SN1, DOPE SN2, DOPG SN1, DOPG SN2, TOCL1 A, TOCL1 B, TOCL1 C, TOCL1 D and surfactant chains. The carbon atom numbers (x-axis) are counted from the carbon atom labelled as shown in Figure S5 for the different molecules. In all cases we find greater disorder induced for the laurate (L20 and L40) systems when compared with the corresponding oleate systems (O20 and O40). Additionally the disorder is greatest toward the membrane mid-plane. In the case of the surfactants greater disorder is observed for the laurate molecules due to the increased hydrophobic mismatch.

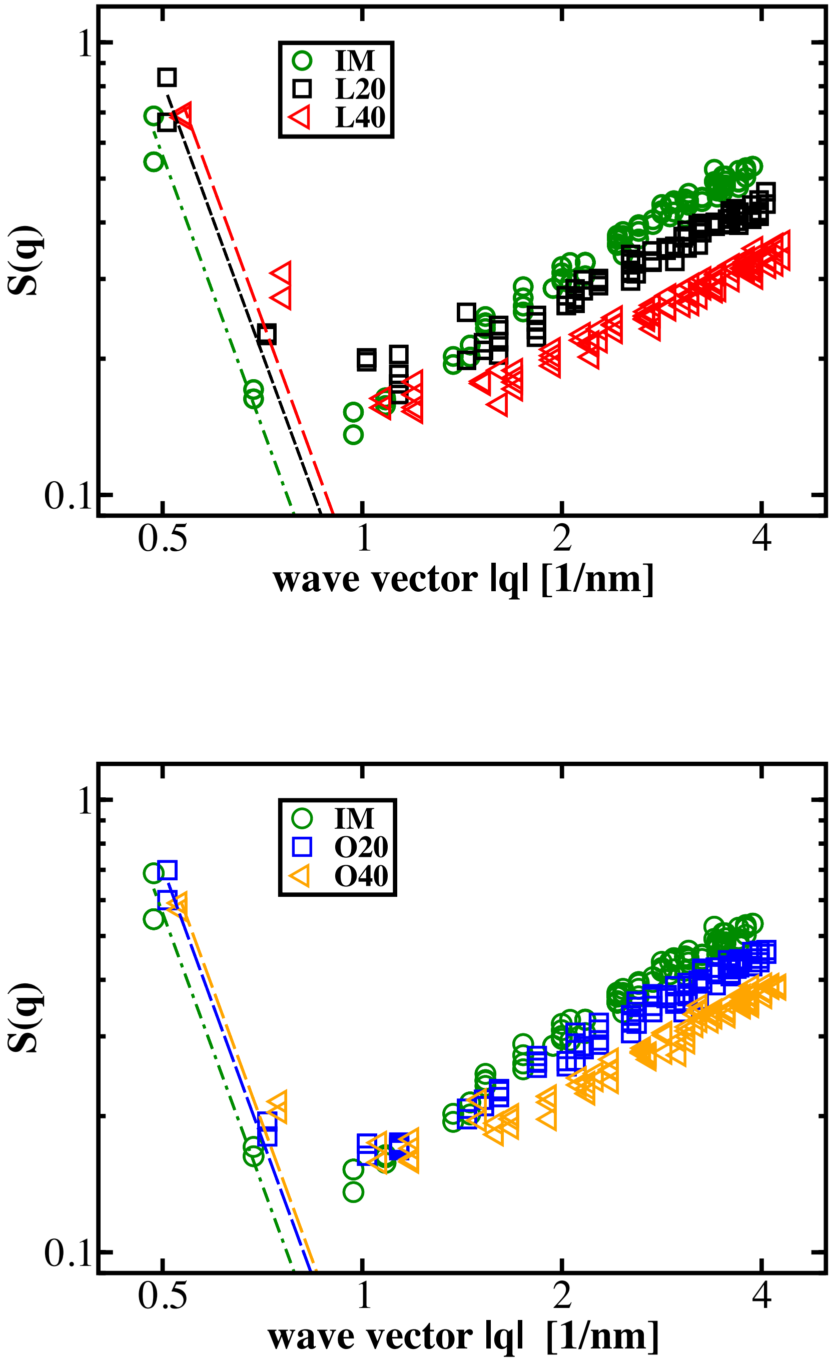

**Fig. S8.** Static structure factor for membrane height fluctuations. The corresponding values of bending modulus are given in main text. Our simulation box sizes are sufficiently large to capture the low $q$ regime in order to extract the bending modulus.

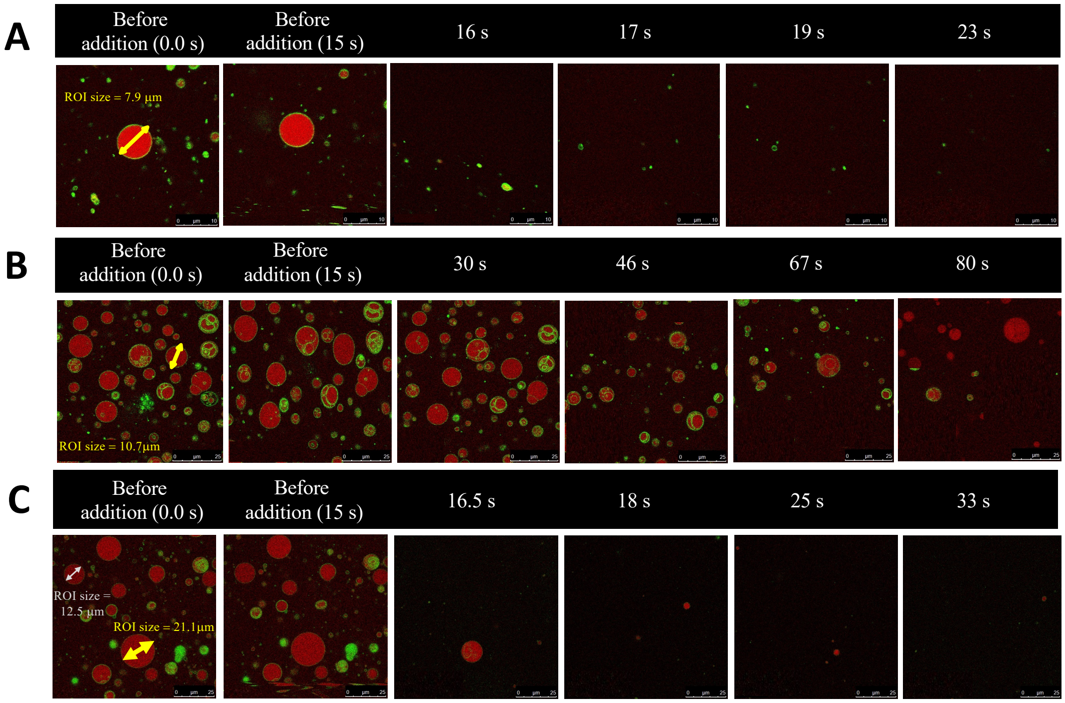

**Fig. S9.** Time-lapse images of multiple GUVs. (A) Addition of sodium laurate of concentration 9 mM (final concentration in the 300 $\mu$l well - GUV size 7.84 $\mu$m) - rupture of GUVs was observed. (B) Addition of sodium oleate of concentration 12 mM (final concentration in the 300 $\mu$l well - GUV size 10.7 $\mu$m) - makes the GUVs to get slowly ruptured. (C) Addition of sodium oleate of concentration 15 mM (final concentration in the 300 $\mu$l well - GUV sizes 12.5 $\mu$m and 21.1 $\mu$m) - makes the GUVs to get ruptured immediately.

### References

1. Agata Witkowska, Lukasz Jablonski, and Reinhard Jahn. A convenient protocol for generating giant unilamellar vesicles containing snare proteins using electroformation. *Scientific reports*, 8(1):1–8, 2018.
2. Pradyumn Sharma, Srividhya Parthasarathi, Nivedita Patil, Morris Waskar, Janhavi S Raut, Mrinalini Puranik, K Ganapathy Ayappa, and Jaydeep Kumar Basu. Assessing barriers for antimicrobial penetration in complex asymmetric bacterial membranes: A case study with thymol. *Langmuir*, 36(30):8800–8814, 2020.
3. Glenn J Martyna, Michael L Klein, and Mark Tuckerman. Nosé–hoover chains: The canonical ensemble via continuous dynamics. *J. Chem. Phys.*, 97(4):2635–2643, 1992.
4. Michele Parrinello and Aneesur Rahman. Polymorphic transitions in single crystals: A new molecular dynamics method. *J. Appl. Phys.*, 52(12):7182–7190, 1981.
5. Tom Darden, Darrin York, and Lee Pedersen. Particle mesh ewald: An nlog (n) method for ewald sums in large systems. *J. Chem. Phys.*, 98(12):10089–10092, 1993.
6. Berk Hess, Henk Bekker, Herman JC Berendsen, and Johannes GEM Fraaije. Lincs: a linear constraint solver for molecular simulations. *J. Comput. Chem*, 18(12):1463–1472, 1997.
7. William Humphrey, Andrew Dalke, Klaus Schulten, et al. Vmd: visual molecular dynamics. *Journal of molecular graphics*, 14(1):33–38, 1996.
8. Ramon Guixà-González, Ismael Rodriguez-Espigares, Juan Manuel Ramírez-Anguita, Pau Carrió-Gaspar, Hector Martinez-Seara, Toni Giorgino, and Jana Selent. Membplugin: studying membrane complexity in vmd. *Bioinformatics*, 30(10): 1478–1480, 2014.
9. Mark James Abraham, Teemu Murtola, Roland Schulz, Szilárd Páll, Jeremy C Smith, Berk Hess, and Erik Lindahl. Gromacs: High performance molecular simulations through multi-level parallelism from laptops to supercomputers. *SoftwareX*, 1:19–25, 2015.
10. W Helfrich. Steric interaction of fluid membranes in multilayer systems. *Zeitschrift für Naturforschung A*, 33(3):305–315, 1978.
